# DLTKcat: deep learning based prediction of temperature dependent enzyme turnover rates

**DOI:** 10.1101/2023.08.10.552798

**Authors:** Sizhe Qiu, Simiao Zhao, Aidong Yang

## Abstract

The enzyme turnover rate, *k*_*cat*_, quantifies enzyme kinetics by indicating the maximum efficiency of enzyme catalysis. Despite its importance, *k*_*cat*_ values remain scarce in databases for most organisms, primarily due to the cost of experimental measurements. To predict *k*_*cat*_ and account for its strong temperature dependence, DLTKcat was developed in this study and demonstrated superior performance (log10-scale RMSE = 0.88, R2 = 0.66) than previously published models. Through two case studies, DLTKcat showed its ability to predict the effect of protein sequence mutations and temperature changes on *k*_*cat*_ values. Although its quantitative accuracy is not high enough yet to model the responses of cellular metabolism to temperature changes, DLTKcat has the potential to eventually become a computational tool to describe the temperature dependence of biological systems.

## 1. Introduction

In the age of synthetic biology, more and more chemical processes are being catalyzed by enzymes [(Stephanopoulos, 2012)][(Madhavan *et al*, 2021)], and therefore, the quantitative study of enzyme kinetics becomes an important topic. The enzyme turnover rate, *k*_*cat*_, is one of the most important parameters in describing enzyme kinetics, which quantifies the maximum efficiency of an enzyme in catalyzing a specific reaction [(Davidi *et al*, 2016)]. In spite of its importance, there currently exists a huge gap of measured *k*_*cat*_ for most organisms in commonly used enzyme databases [(Nilsson *et al*, 2017)], i.e. BRENDA [(Schomburg *et al*, 2017)] and SABIO-RK [(Wittig *et al*, 2018)]. Also, measuring *k*_*cat*_ values via enzyme assays is expensive and labor intensive [(Nilsson *et al*, 2017)], which means that it’s hard to obtain *k*_*cat*_ values in a high-throughput manner. The limited availability of *k*_*cat*_ in databases and the indispensable requirement for *k*_*cat*_ in the study of enzyme kinetics and other fields, such as metabolic modeling [(Adadi *et al*, 2012)], fuels the impetus behind the development of computational methods to predict *k*_*cat*_ values.

There are two main methods to predict *k*_*cat*_ values: 1. estimating *k*_*cat*_ based on apparent catalytic rate (*k*_*app*_) with proteomic and fluxomic profiling; 2. predicting *k*_*cat*_ using the compound protein interaction (CPI) deep learning model. The first method obtains the *k*_*cat*_ value by dividing the measured reaction flux by the quantified protein abundance [(Heckmann *et al*, 2020)][(Bulović *et al*, 2019)]. Although this method has been proved successful in resource allocation models of various microorganisms [(Goelzer *et al*, 2015)][(Jahn *et al*, 2021)][(Coppens *et al*, 2023)][(Wendering *et al*, 2023)], fluxomics and proteomics are costly to measure, making this method difficult to implement.

CPI deep learning models have already been developed to predict biological parameters such as binding affinities (*k*_*d*_) [(Li *et al*, 2020)], michaelis-menten constants (*k*_*m*_) [(Kroll *et al*, 2021)] and enzyme turnover rates (*k*_*cat*_) [(Li *et al*, 2022a)]. The inputs are usually SMILES (simplified molecular-input line-entry system) strings of compounds and subsequences of proteins. Compound and protein features are extracted by graph neural network (GNN), recurrent neural network (RNN) or convolutional neural network (CNN), and then concatenated for the regression of the target value, such as *k*_*cat*_ or *k*_*m*_ [(Lim *et al*, 2021)]. For better performance, attention layers are added to capture the interaction between compound and protein features [(Shin *et al*, 09--10 Aug 2019)][(Karimi *et al*, 2019)]. DLKcat [(Li *et al*, 2022a)], the first CPI deep learning model for *k*_*cat*_ prediction, can predict *log*_10_(*k*_*cat*_) with the RMSE (root mean squared error) score below 1 and Pearson’s r = 0.71 for the test dataset. However, one limitation of DLKcat and most other CPI models is that they do not account for experimental conditions like temperature, pH or ionic strength. As *k*_*at*_ has a strong dependence on temperature [(Arroyo *et al*, 2022)] and temperature is widely available in databases, developing a deep learning model that takes compound, protein and temperature features together as inputs is both necessary and approachable.

In 2023, EF-PreKcat, Revised PreKcat [(Yu *et al*, 2023)] and TurNuP [(Kroll *et al*, 2023)] were developed to predict temperature dependent *k*_*cat*_ values. They all considered *k*_*cat*_ values at different temperatures and include the temperature value as a feature. But none of them assessed the feature importance of temperature or demonstrated any case studies to show the model’s ability to predict the effect of temperature on *k*_*cat*_ values. Also, the R2 (r-squared) scores of predictions by those models were all reported to be below 0.5.

With the aim to construct a deep learning model on *k*_*cat*_ prediction that is more accurate than previously published models, this study developed DLTKcat. DLTKcat is a bi-directional attention CPI model with molecular graphs converted from SMILES strings, 3-mer subsequences of proteins and temperature features as inputs. It showed superior performance (log10-scale RMSE = 0.88, R2 = 0.66) than previously published models (e.g., TurNuP), and demonstrated the feature importance of temperature. Then, DLTKcat exhibited its potential application in enzyme sequence design by predicting the effect of amino acid substitutions on *k*_*cat*_ at different temperatures. Finally, we incorporated temperature dependent proteome constraints in bacterial metabolic modeling with predicted *k*_*cat*_ at different temperatures, to explore the possibility of using DLTKcat to make metabolic modeling sensitive to temperature changes.

## 2. Methods

### 2.1 Dataset preparation

The dataset used to construct the deep learning model was extracted from the BRENDA and SABIO-RK databases. EC number, substrate name, organism name, protein identifier (UniProt ID), enzyme type, temperature and *k*_*cat*_ values were queried from SABIO-RK via application programming interface (API). The data in BRENDA was fetched using BRENDApyrser [(Estévez, 2022)]. The canonical SMILES string [(Weininger, 1988)] of the substrate, that describes the molecular structure of chemical species, was obtained by querying the PubChem compound database [(Kim *et al*, 2023)] via API. The amino acid sequence of each enzyme protein was queried from the UniProt database [(UniProt Consortium, 2023)] based on the UniProt ID also via API. The sequences of wild type (WT) enzymes were mapped directly. For mutants caused by amino acid substitutions, amino acids at mutated locations were changed based on mutation information from BRENDA and SABIO-RK. Entries with other types of mutations were removed. All API codes can be found at https://github.com/SizheQiu/DLTKcat.

After SMILE strings and amino acid sequences were obtained, the dataset filtered out all redundant entries with the same SMILE string, amino acid sequence, temperature and *k*_*cat*_ value. For entries with the same SMILE string, amino acid sequence, temperature but different *k*_*cat*_ values, only the entry with the largest *k*_*cat*_ value was kept, as done in Li et al., 2022 [(Li *et al*, 2022a)]. Finally, 4383 entries from SABIO-RK and 11866 entries from BRENDA remained. 10556 entries’ enzymes were WTs and 5693 entries’ enzymes were mutants (**see SI, Figure S1**). *k*_*cat*_ values of 87 enzyme classes (EC numbers) were found to have significant correlations with temperature, which covered 2430 entries (**see SI, Figure S2**). Considering the uneven distribution of temperature values in the dataset, oversampling was performed to append two times of entries at low (T<20□) and high (T>40□) temperature ranges by randomly duplicating existing entries at those temperature ranges (**see SI, Figure S3**). Because previously published CPI deep learning models have shown that additional features, such as enzyme molar mass or the octanol–water partition coefficient of substrate, could not improve model performance [(Kroll *et al*, 2021)][(Kroll *et al*, 2023)], the finalized dataset of this study only contained SMILES strings of substrates, amino acid sequences of enzyme proteins and temperature values.

### 2.2 Construction of the deep learning model

Similar to other CPI deep learning models, DLTKcat uses Graph Attention Network (GAT) and Convolutional Neural Network (CNN) to extract features from the substrate molecular graph and enzyme protein sequence, respectively (**Figure 1**). The use of bi-directional attention, adopted from BACPI by Li et al., 2022 [(Li *et al*, 2022b)], and integration of temperature and inverse temperature values capture the temperature dependent interactions between atoms of the compound and residues of the protein. Finally, the concatenated features of compound, protein and temperature are fed into several dense layers (fully connected layers) to predict the *log*_10_ *(k*_*cat*_ ) value.

**Figure 1.**
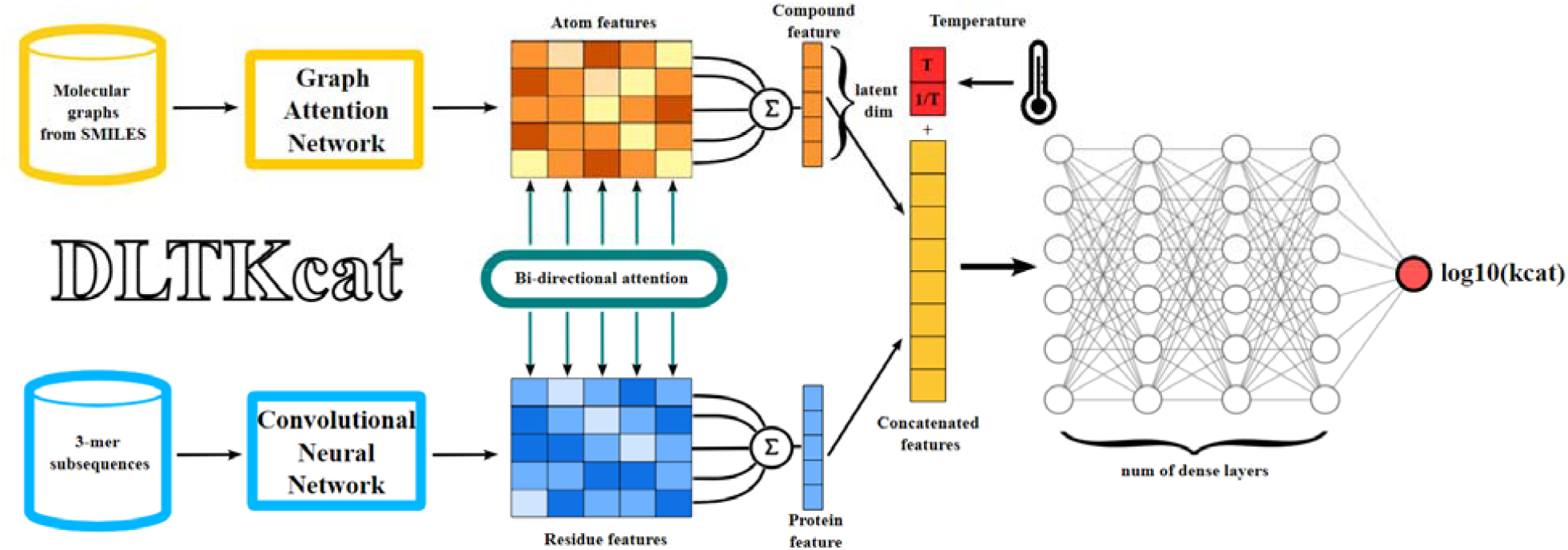
The overview of DLTKcat. With a pair of substrate and enzyme as the input, a GAT and a CNN learn the representations of the atom and residue from the compound molecular graph and protein sequence. Next, atom and residue representations are fed into the bi-directional attention neural network to integrate the representations and capture the important regions of compounds and proteins. Then, temperature () and inverse temperature () are integrated into the concatenated features. Finally, the concatenated features are used to predict the value.

#### 2.2.1 Compound representation

RDKit [(Landrum, 2006)] converts the SMILES string into the molecular graph of the substrate with atoms as vertices, and chemical bonds as edges. The graph, along with the initial embeddings of its vertices, is fed into the graph attentional layer of GAT. A linear learnable transformation converts the embeddings into higher-level features of the compound. The multi-head attention mechanism in GAT concatenates output features from several independent graph attentional layers to increase the stability of the self-attention learning process. Finally, a single-layer neural network transforms concatenated features into the compound space. The final output features are atom features (**Figure 1**). Extended Connectivity Fingerprints (ECFPs) [(Rogers & Hahn, 2010)] of length 1024, computed by RDKit, are also used to represent the compound. A multi-layer neural network transforms ECFPs into the compound space.

#### 2.2.2 Protein representation

To capture diverse protein residue patterns, the protein sequence is split into overlapping 3-mer subsequences. 3-mer subsequences are then translated to randomly initialized embeddings. Through several convolutional layers with leaky ReLU [(Maas *et al*, 2013)] as the activation function, embeddings are transformed to higher-level features of the protein sequence that can capture the complex relationships of residues. The final output features are residue features (**Figure 1**).

#### 2.2.3 Bi-directional attention and integration of temperature

The bi-directional attention mechanism is used to represent the interactions between atoms of the compound and residues of the protein. Residue, atom features and fingerprints are transformed into vectors, and a soft alignment matrix indicates the interaction strengths. The weighted information is extracted from the soft alignment matrix, and attention is computed in both atom-to-residue and residue-to-atom directions. The final outputs are compound and protein features (**Figure 1**). To improve learning stability and representation capacity, a multi-head attention model is used to capture diverse aspects of compound-protein interactions.

Inspired by the Arrhenius equation 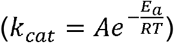 [(Arroyo *et al*, 2022)], temperature (*T*) and inverse temperature 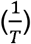 are first normalized 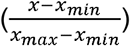, and then concatenated with compound and protein features output by the bi-directional attention process. The inverse of temperature best represents the linear relationship between 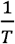 and *log*_10_ *(k*_*cat*_). The concatenated features are then fed into several dense layers, with leaky ReLU as the activation function, for the regression of the *log*_10_ *(k*_*cat*_ ) value.

#### 2.2.4 Model training

Due to the large size of the dataset, batch training was used with a batch size of 32. Adam optimization algorithm [(Kingma & Ba, 2014)] was used to update neural network weights iteratively. The loss function was mean squared error (MSE). The initial learning rate was 0.001, and the learning rate decayed by 50% for every 10 epochs to prevent overfitting. For details of software and hardware, please see the section 1 of supplementary information.

### 2.3 Deep learning model evaluation

The original dataset curated in section 2.1 was randomly split into the test and the train dataset with a ratio of 1:9. The test dataset was held out to examine the accuracy of the model. Before the training, 10% of the train dataset was randomly split as the validation dataset (also called dev set). During the training process, R2 (Eq. 1) and RMSE (Eq. 2) scores of *k*_*cat*_ predictions were computed at each epoch for the test and validation datasets.

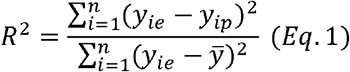

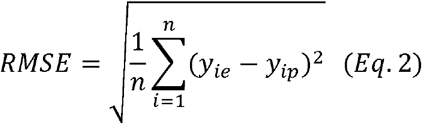

To find the optimal combinations of the latent dimension, the dimension of protein and compound features in bi-directional attention process, and the number of dense layers after feature concatenation (**Figure 1**), training processes were performed for latent dimension = 40, 64 and number of dense layers = 3, 4, 5, 6. The latent dimension and number of dense layers that achieved the lowest RMSE score on the dev set were used to generate the trained deep learning model (**see SI, Figure S4**). The other hyperparameters were the same as default values used in BACPI [(Li *et al*, 2022b)].

1026 entries in the curated dataset had the same protein sequence and substrate as other entries but different temperatures. To assess the feature importance of temperature, they were extracted to obtain R2 and RMSE scores of *k*_*cat*_ predictions with shuffled and unshuffled temperature related features. The feature shuffling was performed by randomly sampling from the original list of feature values, so that the shuffled and unshuffled features still had the same distribution.

### 2.4 Proteome constrained flux balance analysis with predicted *k*_*cat*_

Flux balance analysis (FBA) has been used to estimate metabolic fluxes and cellular growth rates for decades [(Orth *et al*, 2010)]. The basic required inputs are the stoichiometric matrix (*S*) from the genome-scale metabolic model (GSMM) [(Orth *et al*, 2010)] and growth medium parameters that set upper bounds for nutrient uptake rates. FBA computes metabolic fluxes (*v*_*i*_) by maximizing an objective function (Eq. 3), which is usually the growth function (*v*_*growth*_, biomass formation rate normalized to 1 gram dry weight (gDW) of biomass), via linear optimization in a constrained solution space of mass conservation (Eq. 4) and lower/upper bounds (*v*_*lb*_, *v*_*ub*_) of reaction fluxes (Eq. 5). FBA was conducted using COBRApy [(Ebrahim *et al*, 2013)] in this study.

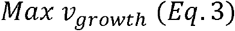

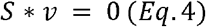

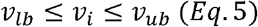

Proteome constrained FBA tightens the solution space by integrating proteome constraints of reactions into conventional FBA [(Mori *et al*, 2016)]. The reaction flux 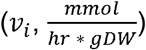 is constrained by the enzyme capacity (*k*_*i*_ [*E*_*i*_] or *a*_*i*_ (*MW*_*i*_*∗* [*E*_*i*_]) (Eq. 6). *k*_*i*_ is the *k*_*cat*_ of reaction *i* [*E*_*i*_] and is the enzyme molar concentration 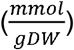. 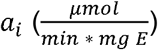 is the enzyme specific activity, defined as the micro moles of products formed by an enzyme in a given amount of time per milligram of the enzyme protein. *MW*_*i*_ is enzyme molar mass 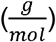. Proteome was divided into sectors of inflexible housekeeping (Q), anabolism (A), transportation (T) and catabolism (C). The upper bound of all flexible sectors (i.e., C, A, T) combined was assumed to be 50% of the total proteome (Eq. 7) [(Zeng & Yang, 2020)][(Regueira *et al*, 2021)][(Qiu *et al*, 2023)].

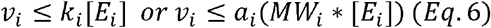

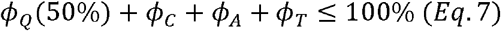

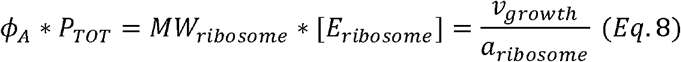

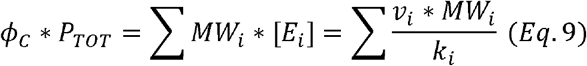

*ϕ*_*x*_ is the mass fraction of sector *x* for *x = A, C, T. P*_*TOT*_ is the total mass of the proteome normalized to 1 gDW of biomass 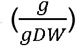. The enzyme activity of the ribosome for the anabolism sector (*a*_*ribosome*_) was set as 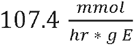 (Eq. 8) [(Schumacher, 2018)][(Regueira *et al*, 2021)]. *k*_*cat*_ values were predicted for the catabolic sector (sector C), by DTLKcat (Eq. 9).

In this study, proteome constrained FBA was performed for *Lactococcus lactis MG1363* (**LL**) and *Streptococcus thermophilus LMG18311* (**ST**). The GSMMs used were obtained from the work of Flahaut et al., 2013 [(Flahaut *et al*, 2013)] and Pastink et al., 2009 [(Pastink *et al*, 2009)]. Experimental data of LL and ST’s growth rates at different temperatures were obtained from Chen et al., 2015 [(Chen *et al*, 2015)] and Vaningelgem et al., 2004 [(Vaningelgem *et al*, 2004)]. The carbon sources of LL and ST, in experiments, were glucose and lactose, respectively. Therefore, the enzyme activities (*a*_*CT*_, CT stands for carbon source transportation) of glucose transport via phosphotransferase system and lactose: galactose antiporter were set as 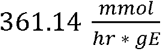 [(Christiansen & Hengstenberg, 1999)], and 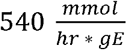 [(Geertsma *et al*, 2005)] (Eq. 10). Because both lactose and glucose were sufficient in the growth medium [(Vaningelgem *et al*, 2004)][(Chen *et al*, 2015)], no michaelis-menten kinetics was needed for transporter proteins. Lactic and acetic acids were two major products of the central carbon metabolism of lactic acid bacteria, and the enzyme activity of acid exportation (*a*_*AT*_) was set as 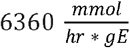 [(Schumacher, 2018)][(Regueira *et al*, 2021)] (Eq. 10).

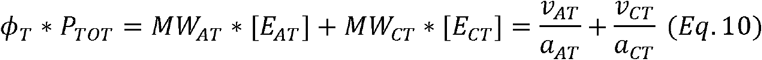

Temperature dependent *k*_*cat*_values were predicted for enzymes in two bacteria’s central carbon metabolism (**see SI, Table S1 and S2**). The SMILES strings of substrates were queried from PubChem with metabolite names in GSMMs, and protein sequences were queried from UniProt with gene locus tags in genome assemblies of LL, GCF_000009425.1 [(Wegmann *et al*, 2007)], and ST, GCF_000011825.1 [(Bolotin *et al*, 2004)]. The predicted *k*_*cat*_ for the primary substrate of each reaction was selected as the *k*_*cat*_ of the reaction. For isozymes that catalyze the same metabolic reaction, the largest *k*_*cat*_ was selected. Both ST and LL are important and widely used lactic acid bacteria, but their enzyme *k*_*cat*_ values are quite limited in databases. For example, there are only 11 entries for ST in SABIO-RK, most were contributed by Simon and Hofer, 1981 [(Simon & Hofer, 1981)]. Therefore, this study used DLTKcat to fill the gap and examined DLTKcat’s performance in predicting metabolic responses to temperature changes.

## 3. Results

### 3.1 DLTKcat outperforms previous models on temperature dependent *k*_*cat*_ prediction

With optimal hyperparameters (**see SI, Figure S4**), the model training process reduced RMSE scores of predicted *log*_10_ *(k*_*cat*_ ) of the test dataset from 1.33 to 0.88, and enhanced R2 scores from 0.25 to 0.66 after 20 epochs (**Figure 2A**). The R2 scores of previously published deep learning models on temperature dependent were all reported to be below 0.5 [(Yu *et al*, 2023)][(Kroll *et al*, 2023)], and DLTKcat has outperformed them by reaching a R2 score of 0.66 (**Figure 2B**). In addition, DLTKcat showed good prediction accuracy with low RMSE scores for datasets of experimental values at the lower 25%, middle 50% and upper 25% ranges (**Figure 2C**). In a nutshell, DLTKcat demonstrated superior performance in comparison to previously published deep learning models, and a robust accuracy for target values (experimental values) at different ranges.

**Figure 2.**
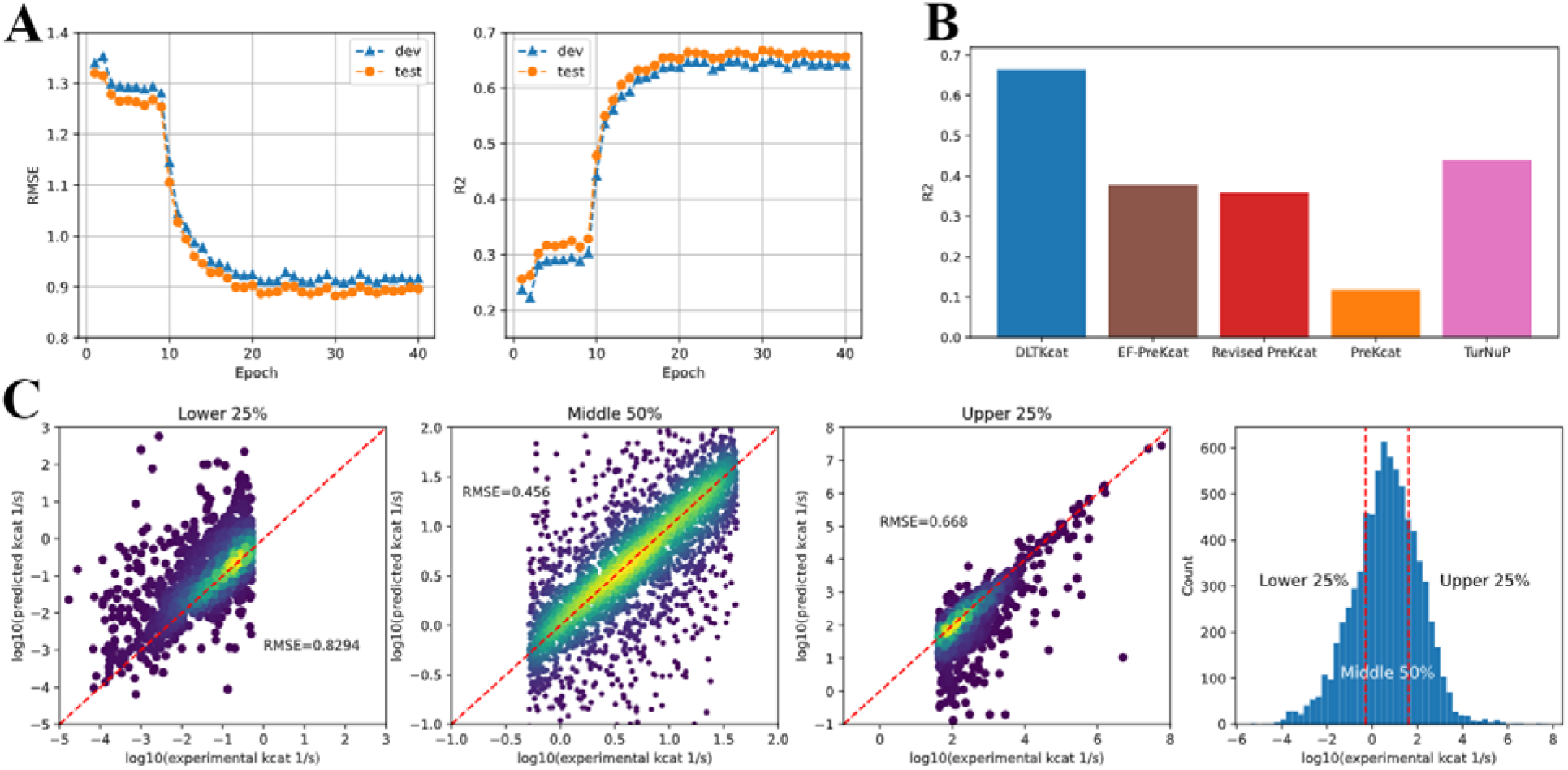
Assessment of the model performance. (A) The RMSE and R2 scores of prediction during the training process. test: the test set; dev: the validation set. (B) Comparison of R2 scores of DLTKcat, EF-PreKcat, Revised PreKcat, PreKcat [(Yu *et al*, 2023)] and TurNuP [(Kroll *et al*, 2023)] on prediction with temperature values. (C) RMSE scores of prediction for datasets with experimental values at lower 25%, middle 50% and upper 25%.

Subsequently, the accuracy of DLTKcat for WT/mutant enzymes and enzymes from different metabolic pathways was examined. For both WTs and mutants, the R2 and RMSE scores were all around 0.8 and 0.6 respectively (**Figure 3A**). When predicting values for different metabolic pathways, DLTKcat showed its ability to discriminate enzymes in primary metabolism - catabolism/energy (primary-CE) and other pathways, with a significant higher distribution (p-value < 0.05) of predicted values in primary-CE (**Figure 3B**). Also, DLTKcat could predict values for enzymes from different metabolic pathways with R2 scores around 0.8 and RMSE scores around 0.6. In short, DLTKcat could well characterize enzymes from different contexts.

**Figure 3.**
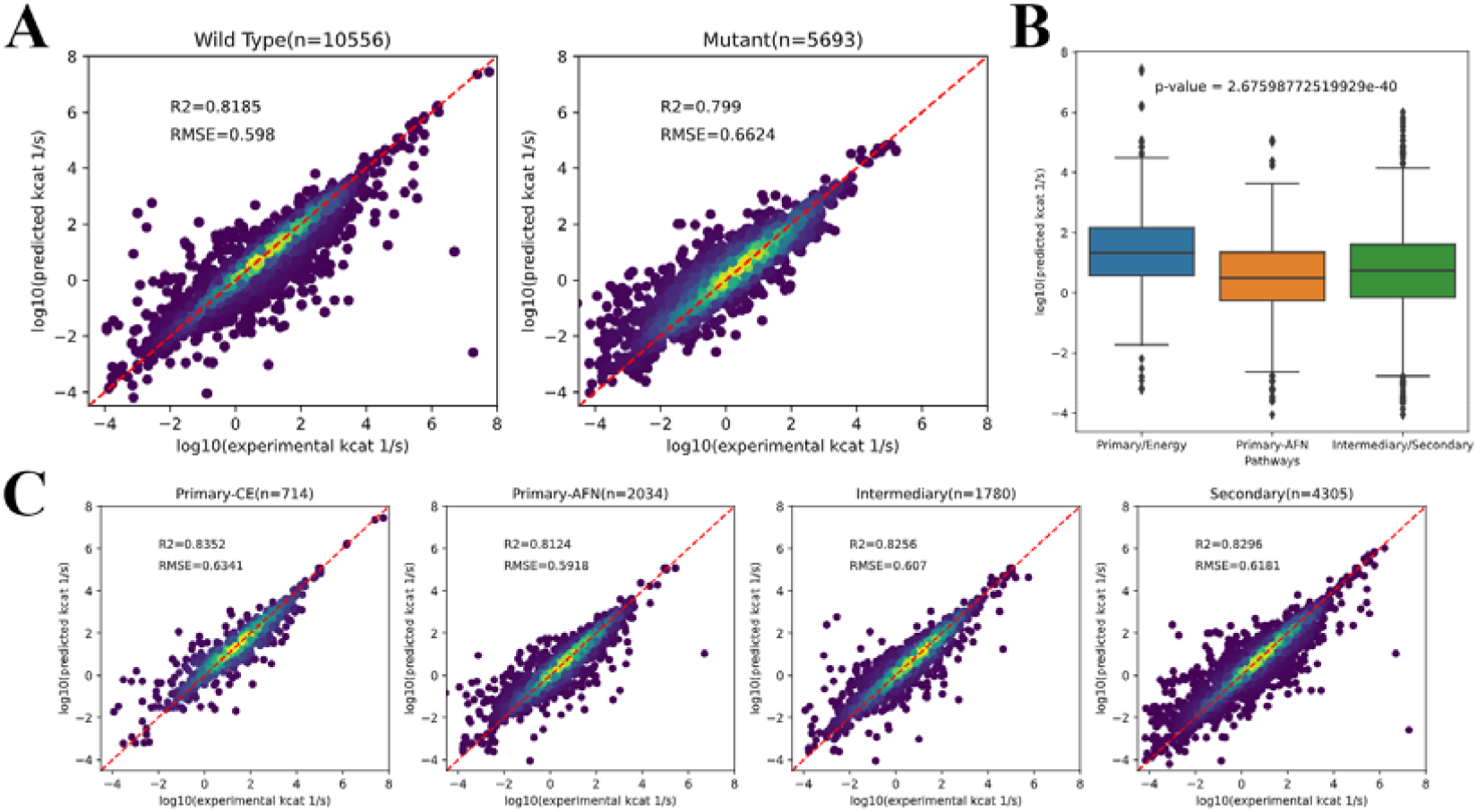
The accuracy of prediction for enzymes from different contexts. (A) R2 and RMSE scores of prediction for WT and mutant enzymes. (B) The comparison of distributions of predicted values in different metabolic pathways (p-value < 0.05). (C) R2 and RMSE scores of prediction for enzymes in primary metabolism - catabolism/energy (primary-CE), primary metabolism - amino acid/fatty acid/nucleotide (primary-AFN), intermediary and secondary metabolism.

### 3.2 The contribution of temperature related features to prediction

Before feature importance analysis, the prediction accuracy was examined for different reflected temperature ranges (*T* < 20°C ≤ *T* ≤ 40°C and *T* > 40°C). High R2 and low RMSE scores reflected that DLTKcat could accurately predict *k*_*cat*_ for low, middle and high temperatures, with an error far below one order of magnitude (**Figure 4A**). Then, feature shuffling, also known as feature permutation, was performed to show the importance of temperature and inverse temperature values. It was observed that the RMSE score increased by around 0.1 and R2 score decreased by around 0.1 when temperature related features were shuffled (**Figure 4B**). For high (*T* > 40°C) and low (*T* < 20°C) temperature ranges, the increase of RMSE and decrease of R2, caused by feature shuffling, became larger (**Figure 4CD**). In short, the decrease in prediction accuracy with shuffled temperature related features demonstrated the importance of temperature related features in DLTKcat.

**Figure 4.**
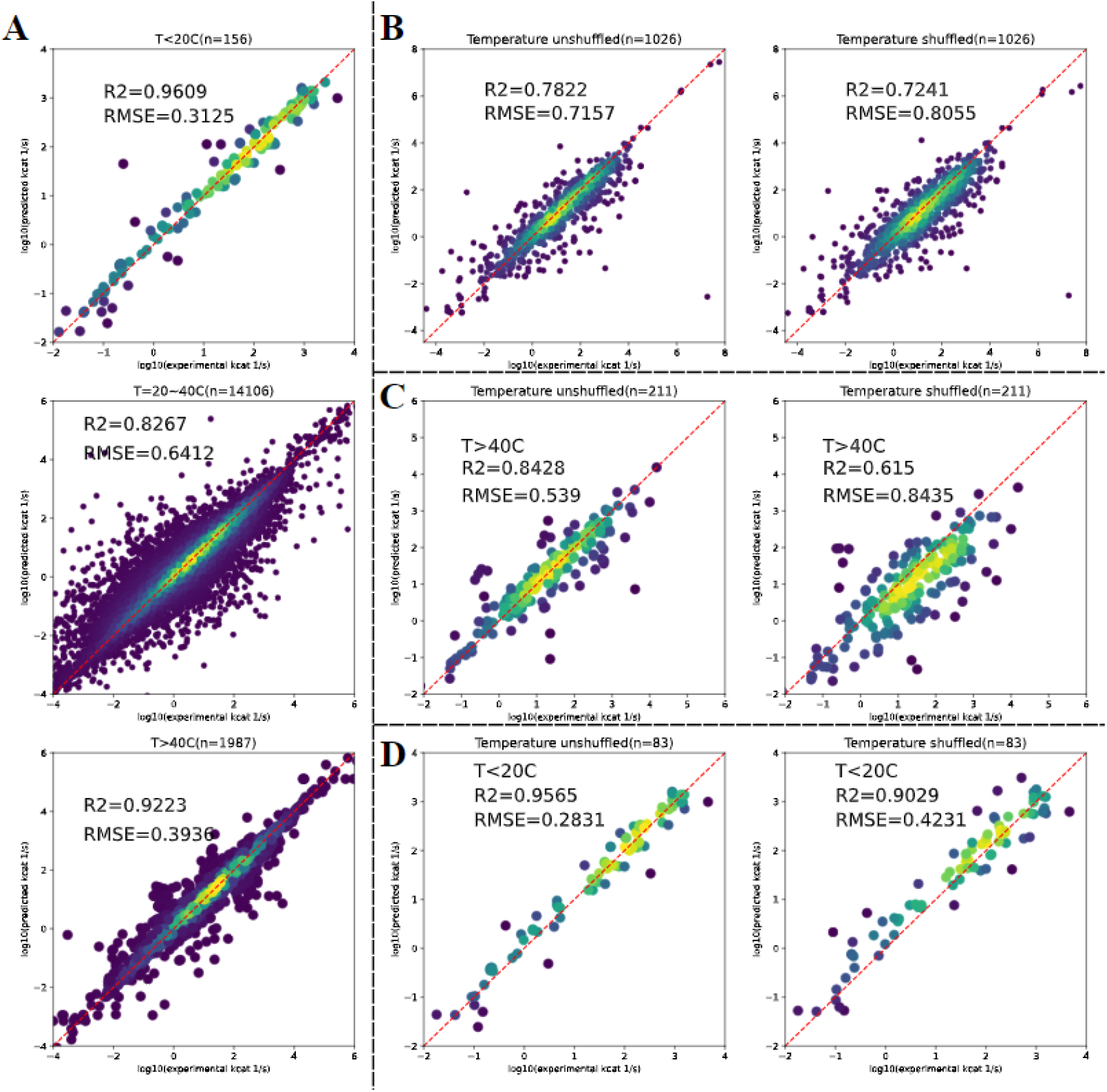
The importance of temperature related features in DLTKcat. (A) R2 and RMSE scores of predicted values at low (<20□), middle (20∼40□) and high (>40□) temperature ranges. (B) R2 and RMSE scores of predicted values with unshuffled and shuffled temperature related features for the selected dataset with 1026 entries. (C) R2 and RMSE scores of predicted values with unshuffled and shuffled temperature related features for entries of high temperature. (D) R2 and RMSE scores of predicted values with unshuffled and shuffled temperature related features for entries of low temperature.

### 3.3 DLTKcat can predict the effect of protein sequence mutations on

*Pyrococcus furiosus* Ornithine Carbamoyltransferase values of WT and mutants, generated via directed evolution, at 30□ and 55□were obtained from Roovers et al., 2001 [(Roovers *et al*, 2001)]. The protein sequence of *Pyrococcus furiosus* Ornithine Carbamoyltransferase was obtained from Uniprot with the Uniprot ID of Q51742. The prediction achieved high accuracy (RMSE = 0.5) (**Figure 5A**). Predicted values at 55□ were higher than those at 30□ (**Figure 5ABC**), which was both consistent with the experimental data and the nature of *Pyrococcus furiosus* being a hyperthermophile favoring high temperature [(Fiala & Stetter, 1986)].

**Figure 5.**
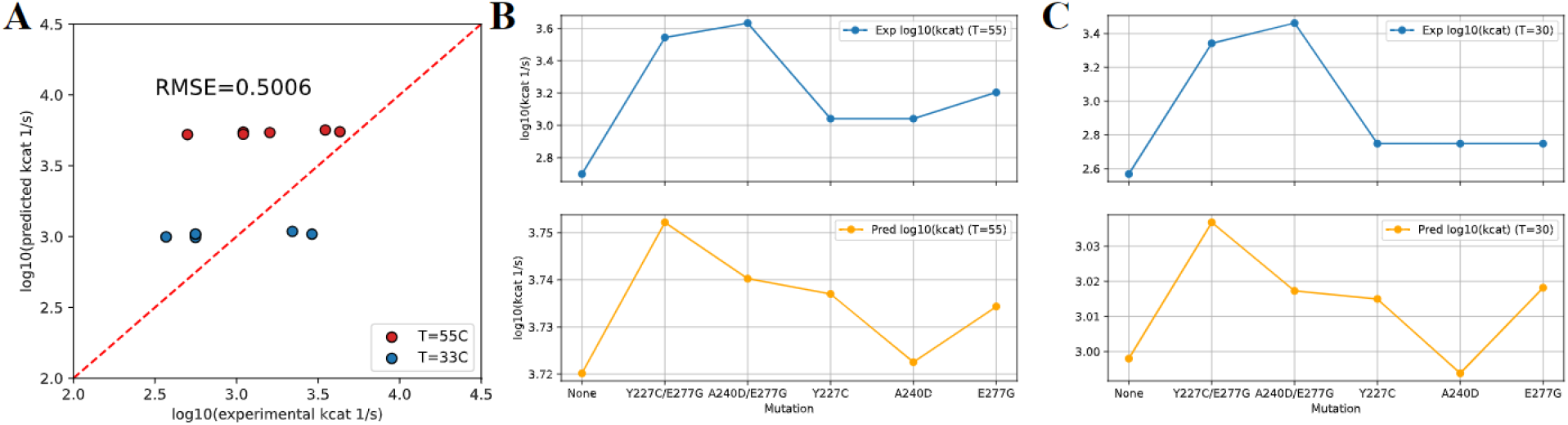
Prediction of the effect of SAVs on values. (A) Comparison between experimental and predicted of Pyrococcus furiosus Ornithine Carbamoyltransferase, RMSE = 0.5006. (B) Experimental (blue line) and predicted (orange line) values of WT and mutants at 55□. (C) Experimental (blue line) and predicted (orange line) *log*_10_ *(k*_*cat*_ ) values of WT and mutants at 30□. Exp: experimental value; Pred: predicted value.

With respect to the effect of mutations, DLTKcat suggested that amino acid substitutions at 227th, 240th and 277th amino acids could increase the value, consistent with the experimental data, despite that the numerical difference between predicted values of mutants and WT was small (**Figure 5BC**; note the difference in scale between upper and lower y-axes). Furthermore, DLTKcat also captured that the combination of two amino acid substitutions, Y227C/E277G and A240D/E277G, could result in greater improvement on the value than the substitution at each single site, though it failed to predict that the of A240D/E277G was higher than that of Y227C/E277G (**Figure 5BC**).

### 3.4 Temperature sensitive metabolic modeling with predicted *k*_*cat*_

DLTKcat predicted *k*_*cat*_ values for enzymes of *Lactococcus lactis MG1363* (LL) at 30, 32, 34, 36 and 38 □, and of *Streptococcus thermophilus LMG18311* (ST) at 25, 32, 37, 42, 46 and 49 □, which were temperatures where LL and ST’s growth rates were measured in experimental data [(Vaningelgem *et al*, 2004)], [(Chen *et al*, 2015)]. DLTKcat predicted that *k*_*cat*_ of most catabolic enzymes in LL would decrease when temperature increased from 30 □ to 38 □, especially for glucose-6-phosphate isomerase (PGI), phosphofructokinase (PFK), phosphoglycerate kinase (PGK), pyruvate kinase (PYK), pyruvate formate lyase (PFL) and phosphotransacetylase (PTAr) (**Figure 6A**). The predicted decrease of the activity of catabolism in LL in response to temperature increase is consistent with the experimental observation that LL stopped growing after temperature became larger than 38 □ [(Chen *et al*, 2015)]. For catabolic enzymes in ST, DLTKcat predicted that most enzymes’ *k*_*cat*_ would increase when temperature increased from 25 □ to 42 □, especially for fructose-bisphosphate aldolase (FBA), Glyceraldehyde-3-phosphate dehydrogenase (GAPD), phosphoglycerate mutase (PGM), enolase (ENO) and pyruvate kinase (PYK) (**Figure 6B**). The predicted increase of catabolic activity in ST when temperature increases to 42 □ is consistent with both the experimental data [(Vaningelgem *et al*, 2004)] and the nature of ST being a thermophile [(Harnett *et al*, 2011)]. These results showed that, in general, DLTKcat could qualitatively predict metabolic responses of bacteria to certain temperature changes.

**Figure 6.**
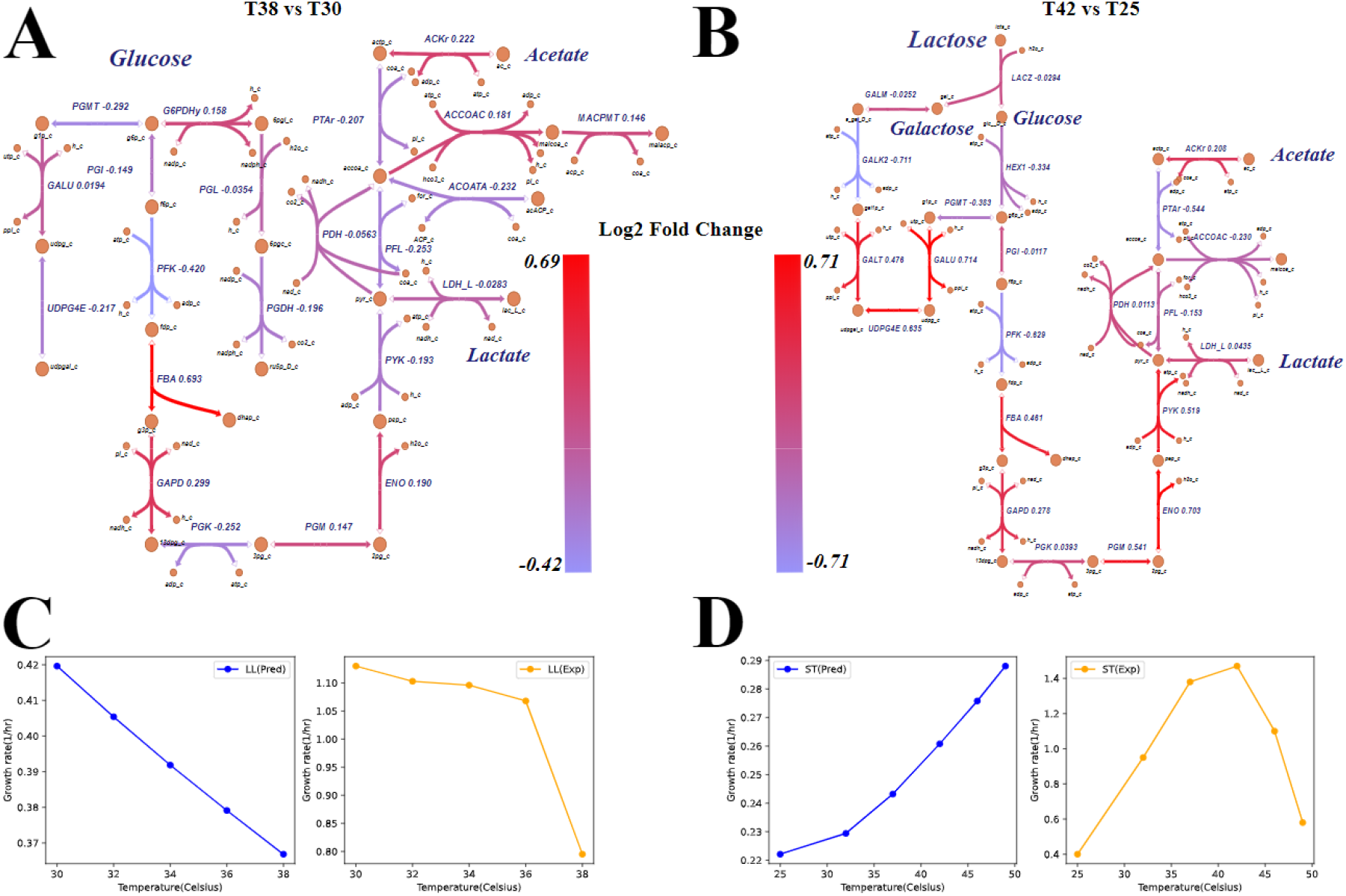
Prediction of bacteria metabolism at different temperatures. (A) Log2 fold change of predicted *k*_*cat*_ values for LL at 38 □ and 30 □ (38 □ vs 30 □). (B) Log2 fold change of predicted *k*_*cat*_ values for ST at 42 □ and 25 □ (42 □ vs 25 □). (C) Comparison of predicted (blue) and experimental (orange) growth rates of LL at 30, 32, 34, 36 and 38 □. (D) Comparison of predicted (blue) and experimental (orange) growth rates of ST at 25, 32, 37, 42, 46 and 49 □. LL: *Lactococcus lactis MG1363*; ST: *Streptococcus thermophilus LMG18311*; Exp: experimental value; Pred: predicted value. Reaction information can be found in **SI, Table S1&2**.

However, the quantitative accuracy of growth rates computed by proteome constrained FBA was low. In proteome constrained FBA for LL, the *k*_*cat*_ of fructose-bisphosphate aldolase (FBA) in LL was fixed at -[(Callens *et al*, 1991)] in sacrifice of temperature sensitivity, because predicted values at different temperatures were unrealistically low (-), compared with experiment values in other bacteria [(Callens *et al*, 1991)][(Plater *et al*, 1999)]. The predicted growth rates of LL by proteome constrained FBA captured the decreasing trend in response to the increase of temperature, but the predicted values were deviant from experimental values (**Figure 6C**). The proteome constrained FBA predicted the increase of ST&#x2019;s growth rate from 25&□to 42&□, but it failed to predict the drop of growth rate from 42 &□ to 49 &□ (**Figure 6D**). Also, the predicted increase of values from 42 &□ to 49 &□ by DLTKcat (**see SI, Figure S5**) contradicted the experimental finding that 49 &□ is close to the theoretical maximum temperature for ST to survive, 47&#x223C;50 &□ [(Harnett *et al*, 2011)]. To conclude, the log10-scale RMSE score within 1 of DLTKcat is not low enough to enable temperature sensitive proteome constraints to enable proteome constrained FBA to predict bacterial growth and metabolism with good quantitative accuracy.

## 4. Discussion

The expensive cost of obtaining enzyme *k*_*cat*_ values in wet lab stimulates the need of developing computational models to predict *k*_*cat*_ . Nevertheless, predicting temperature dependent *k*_*cat*_ is a challenging task, as temperature is not only a variable in the exponential factor of the Arrhenius equation, it also affects the activation energy of the enzyme catalyzed reaction, which is governed by the compound protein interaction [(Arroyo *et al*, 2022)]. To tackle the challenging task, this study constructed a CPI deep learning model called DLTKcat. DLTKcat used bi-directional attention mechanism, whose advantage in capturing important regions in compounds and proteins was already shown by BACPI [(Li *et al*, 2022b)]. The use of both temperature and inverse temperature values facilitated the learning process of the neural network by representing features in the most biophysical relevant form to *k*_*cat*_ [(Arroyo *et al*, 2022)]. Also, oversampling on entries at low and high temperature ranges compensated for the imbalanced distribution of temperature in the dataset (**see SI, Figure S3**). As a result, DLTKcat showed superior performance (log10-scale RMSE = 0.88, R2 = 0.66) than previously published models (e.g., Revised PreKcat, TurNuP) and robust accuracy for *k*_*cat*_ predictions in different contexts. In addition, feature shuffling demonstrated the contribution of temperature related features to this deep learning model.

By accurately predicting the effect of protein sequence mutations on the *k*_*cat*_ value of *Pyrococcus furiosus* Ornithine Carbamoyltransferase at different temperatures (**section 3.3**), DLKcat exhibited its function in scoring the efficiency of in-silico designed enzyme protein sequences. Imaginably, the combination of DLTKcat and optimization algorithms (e.g., genetic programming) can become a computational tool to design site-specific mutagenesis to optimize enzyme catalysis, which will be more efficient than directed evolution that relies on random mutagenesis.

Nonetheless, the second case study (**section 3.4**) of generating temperature dependent proteome constraints for metabolic modeling revealed the limitation of DLTKcat that its prediction error was not low enough to accurately model the response of cellular metabolism to temperature changes. Because all *k*_*cat*_ values of catabolic enzymes in ST and LL were predicted by DLTKcat, the propagation of error led to the inaccuracy of proteome constrained FBA. In short, deep learning can gap fill a few missing *k*_*cat*_ values in the metabolic network, as done in Li et al., 2022 [(Li *et al*, 2022a)], but the accuracy of proteome constrained FBA will not be high if most proteome constraints are based on predicted *k*_*cat*_values.

To further improve the performance and utility of DLTKcat, including additional experimental conditions like pH, metal ion concentrations might be an approach, but the lack of data restricted existing models from accounting for those factors [(Yu *et al*, 2023)]. Including the optimal enzyme temperature either from databases or predictions [(Gado *et al*, 2020)] might be able to enhance the temperature sensitivity of DLTKcat. The difference between the of experimental temperature and optimal temperature could inform the model whether the temperature feature has a negative or positive effect on the *k*_*cat*_ value. However, the success this approach depends on the accuracy of enzyme optimal temperature prediction, which was reported to have a RMSE around 2 [(Li *et al*, 2019)][(Gado *et al*, 2020)].

Overall, DLTKcat can provide accurate predictions of *k*_*cat*_ and account for the effect of temperature changes. Two case studies (3.3 and 3.4) have revealed potential applications of DLTKcat on protein engineering, bacterial phenotype prediction, etc. Additionally, DLTKcat can be easily modified to predict other temperature dependent CPIs, such as [(Quinlan, 1980)][(Maggi *et al*, 2018)]. With future improvements of the model framework, DLTKcat, as we envisage, will become a computational tool to quantitatively model the temperature dependence of biological systems, and contribute to the development of bioprocess digital twins.

## Abbreviation

API: application programming interface
CNN: convolutional neural network
CPI: compound protein interaction
ECFP: Extended Connectivity Fingerprint
GAT: graph attention network
gDW: gram dry weight
GNN: graph neural network
LL: *Lactococcus lactis MG1363*
Leaky ReLU: leaky Rectified Linear Unit
MSE: mean squared error
R2: r-squared, the coefficient of determination
RMSE: root mean squared error
RNN: recurrent neural network
SAV: single amino acid variation
SMILES: simplified molecular-input line-entry system
ST: *Streptococcus thermophilus LMG18311*
WT: wild type

## Acknowledgements

The authors would like to acknowledge the use of the University of Oxford Advanced Research Computing (ARC) facility (http://dx.doi.org/10.5281/zenodo.22558) in carrying out this work. The authors would like to thank Yichi Zhang for offering technical guidance.

## Author contributions

Sizhe Qiu constructed the deep learning model and performed case studies. Simiao Zhao assisted in the construction of the deep learning model. Aidong Yang supervised this research project and critically reviewed the manuscript.

## Data availability statement

The code and data are openly available at https://github.com/SizheQiu/DLTKcat.

