## Supplemental Table S1 and S2, Figure S1, S2, S3, S4 and S5 for "DLTKcat: deep learning based prediction of temperature dependent enzyme turnover rates"

### Authorship

Sizhe Qiu^1^, Simiao Zhao^2^, Aidong Yang^1^*

^1^Department of Engineering Science, University of Oxford, OX1 3PJ, United Kingdom

^2^Radcliffe Department of Medicine, University of Oxford, OX3 9DU, United Kingdom

### 1. Software and code availability

All scripts were written in python. The deep learning model was implemented using PyTorch v1.7.1. The computer used in this work was a Dell Latitude Laptop with intel core i7 CPU. The model was trained with GPU RTX8000 provided by Advanced Research Computing (ARC) service in the University of Oxford [(Richards, 2015)]. Figures were edited using InkScape (<https://inkscape.org/>). The code and data used to generate results of this paper are available at <https://github.com/SizheQiu/DLTKcat>.

### 2. Tables

| Table S1. Metabolic reaction information of *Lactococcus lactis MG1363* | | |
| --- | --- | --- |
| ID | name | EC number |
| GLCpts | D-glucose transport via PEP:Pyr phosphotransferase system | _ |
| G6PDH | Glucose-6-phosphate dehydrogenase | 1.1.1.49 |
| PGL | 6-phosphogluconolactonase | 3.1.1.31 |
| PGDH | 6-phosphogluconate dehydrogenase | 1.1.1.351, 1.1.1.44 |
| GALU | UTP-glucose-1-phosphate uridylyltransferase | 2.7.7.9 |
| PGMT | Phosphoglucomutase | 5.4.2.2, 5.4.2.5 |
| UDPG4E | UDPglucose 4-epimerase | 5.1.3.2 |
| PGI | Glucose-6-phosphate isomerase | 5.3.1.9 |
| PFK | Phosphofructokinase | 2.7.1.11 |
| FBA | Fructose-bisphosphate aldolase | 4.1.2.13 |
| TPI | Triose-phosphate isomerase | 5.3.1.1 |
| GAPD | Glyceraldehyde-3-phosphate dehydrogenase | 1.2.1.12 |
| PGK | Phosphoglycerate kinase | 2.7.2.3 |
| PGM | Phosphoglycerate mutase | 5.4.2.11 |
| ENO | Enolase | 4.2.1.11 |
| PYK | Pyruvate kinase | 2.7.1.40 |
| LDH | Lactate dehydrogenase | 1.1.1.27 |
| PFL | Pyruvate formate lyase | 2.3.1.54 |
| PDH | Pyruvate dehydrogenase | 1.2.7.1 |
| PTAr | Phosphotransacetylase | 2.3.1.8 |
| ACKr | Acetate kinase | 2.7.2.1 |
| ACCOAC | Acetyl-CoA carboxylase | 6.4.1.2 |
| MACPMT | Malonyl CoAacyl carrier protein S malonyltransferase | 2.3.1.39 |

| Table S2. Metabolic reaction information of *Streptococcus thermophilus LMG18311* | | |
| --- | --- | --- |
| ID | name | EC number |
| LCTSGALex | Lactose galactose exchange via antiporter | _ |
| LACZ | Beta-galactosidase | 3.2.1.23 |
| GALK | Galactokinase | 2.7.1.6 |
| HEX | Hexokinase (D-glucose:ATP) | 2.7.1.2 |
| GALK | Galactokinase | 2.7.1.6 |
| GALM | Aldose 1-epimerase | 5.1.3.3 |
| GALT | Galactose 1 phosphate uridylyltransferase | 2.7.7.10 |
| GALU | UTP-glucose-1-phosphate uridylyltransferase | 2.7.7.9 |
| PGMT | Phosphoglucomutase | 5.4.2.2, 5.4.2.5 |
| UDPG4E | UDPglucose 4-epimerase | 5.1.3.2 |
| PGI | Glucose-6-phosphate isomerase | 5.3.1.9 |
| PFK | Phosphofructokinase | 2.7.1.11 |
| FBA | Fructose-bisphosphate aldolase | 4.1.2.13 |
| TPI | Triose-phosphate isomerase | 5.3.1.1 |
| GAPD | Glyceraldehyde-3-phosphate dehydrogenase | 1.2.1.12 |
| PGK | Phosphoglycerate kinase | 2.7.2.3 |
| PGM | Phosphoglycerate mutase | 5.4.2.11 |
| ENO | Enolase | 4.2.1.11 |
| PYK | Pyruvate kinase | 2.7.1.40 |
| LDH | Lactate dehydrogenase | 1.1.1.27 |
| PFL | Pyruvate formate lyase | 2.3.1.54 |
| PDH | Pyruvate dehydrogenase | 1.2.7.1 |
| PTAr | Phosphotransacetylase | 2.3.1.8 |
| ACKr | Acetate kinase | 2.7.2.1 |
| ACCOAC | Acetyl-CoA carboxylase | 6.4.1.2 |

Reaction IDs, names, and enzyme EC numbers are all obtained from the BIGG database (<http://bigg.ucsd.edu/>) [(King *et al*, 2016)].

### 3. Figures

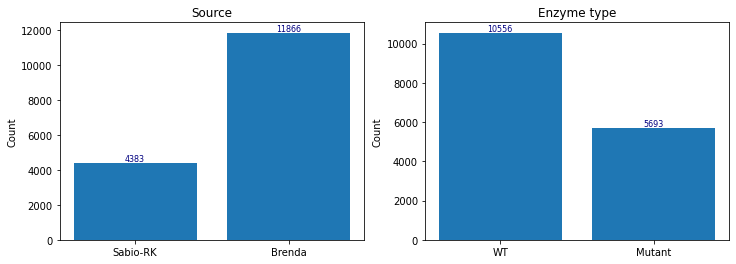

Figure S1. statistics of collected data from enzyme databases. Left: 4383 entries are from Sabio-RK, 11866 entries are from Brenda. Right: enzymes in 10556 entries are wild types, in 5693 entries are mutants.

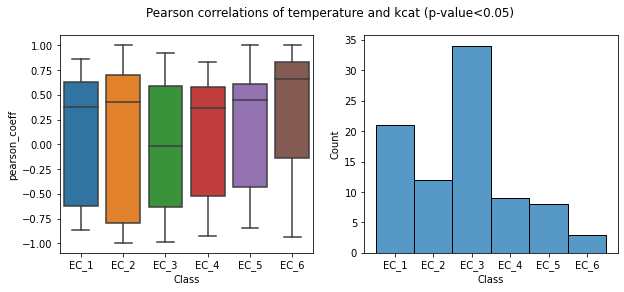

Figure S2. Significant Pearson correlations of $k_{cat}$ of 87 enzyme classes (EC numbers) covering 2430 entries and temperature.

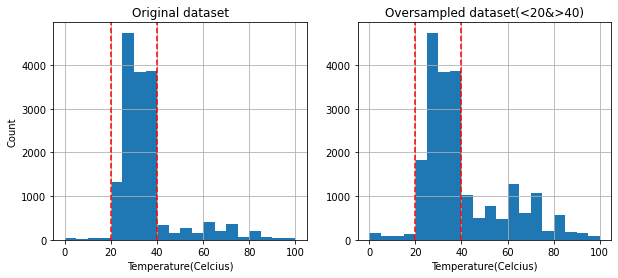

Figure S3. Training datasets before and after oversampling of entries at low and high temperature ranges.

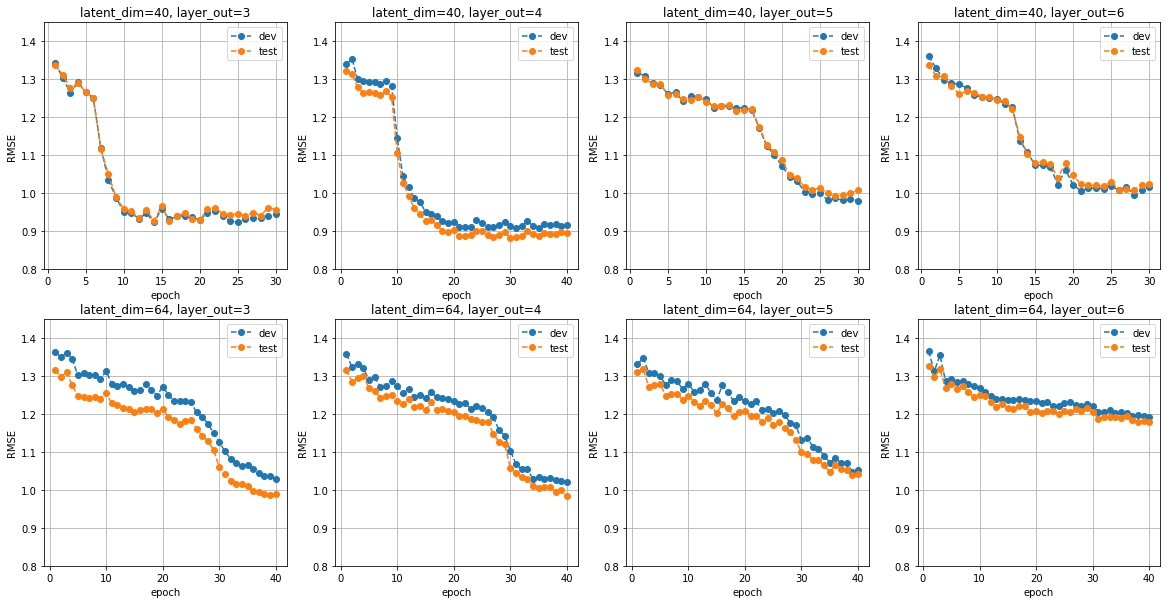

Figure S4. Results of hyperparameter optimization for latent dimension (latent_dim) and number of dense layers (layer_out). The trial with latent_dim=40 and layer_out=4 achieved the lowest RMSE. test: the test set; dev: the validation set (the development set).

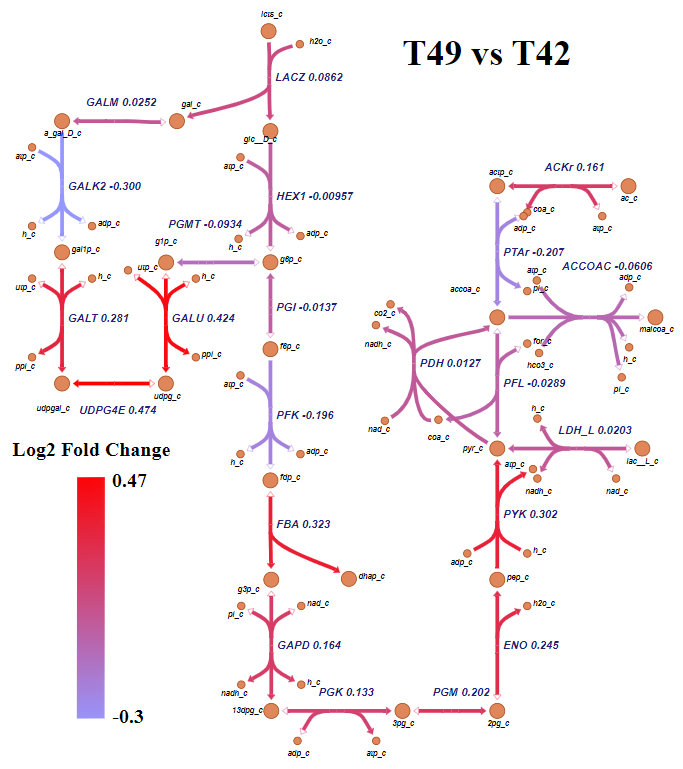

Figure S5. Log 2 fold change of predicted $k_{cat}$ values for ST at 49 ℃ and 42 ℃ (49 ℃ vs 42 ℃).
